# Shoal cohesion and polarisation increase in turbid water and improve foraging performance

**DOI:** 10.1101/2025.04.28.650972

**Authors:** Christos C. Ioannou, James Cordery, Sean A. Rands

**Author notes:** Corresponding author: Christos C. Ioannou.

## Abstract

Group living can make animals more resilient or more susceptible to human-induced rapid environmental change, and whether animals demonstrate plasticity in their collective behaviour to mitigate the effects of environmental change is unknown. Freshwater and coastal habitats are becoming increasingly turbid as a result of human activity, which limits the availability of information and hence responses to critical stimuli such as food and predators. Here, we tested whether the well-documented improvement in foraging performance by larger groups is affected by moderate levels of water turbidity. Fish shoals were faster to attack the food stimulus in larger groups and slower in turbid water, effects which were independent of one another. However, shoals became more cohesive and coordinated (i.e. more polarised in direction) in turbid water, and this corresponded to more cohesive and polarised shoals being faster to attack the food stimulus in turbid water but slower in clear water. Our results demonstrate that fish shoals can increase cohesion and coordination in moderate levels of turbidity, that this plasticity improves their response to ecologically relevant ephemeral stimuli, and that collective behaviour, rather than group size, can mediate the response of groups to environmental change.

## Introduction

Human-induced rapid environmental change is affecting the way that animals sense and respond to ecologically critical stimuli such as food and predators [1–3]. Some of these effects are through changes to an organism’s surroundings that increase environmental noise, where such noise is defined as environmental conditions that can “interfere with the sensation of biologically meaningful stimuli” [4]. Acoustic noise from road or boat traffic, for example, can reduce foraging efficiency [5] and delay anti-predator responses [6]. However, an advantage for animals that live in groups is an enhanced ability to detect and process information, resulting in improved decision making that is widely referred to as collective or swarm intelligence [7,8]. Group living may thus make animals more robust to changes in their abiotic environment, mitigating some of the costs of anthropogenic change.

Social interactions can, however, make animals more susceptible to shifting environmental conditions, for example where adaptation is delayed by a reliance on outdated, socially-learnt information [9]. An increased susceptibility in group living animals can also occur because the environmental noise that interferes with the sensation of stimuli also disrupts the social interactions that determine collective behaviour [10,11]. In situations where a group’s collective behaviour is critical in responding to stimuli [12,13], then the negative impacts of the environmental noise on responses is indirect, occurring via reduced cohesion and coordination.

Here, we focus on water turbidity which is increasing in freshwater and coastal habitats across the globe as a result of human activity including agriculture, quarrying and development of infrastructure [14,15]. Turbidity is created by the scattering of light by particles suspended in water, and acts as environmental noise by reducing the distance over which stimuli can be detected visually [16–19]. Such a constraint on visual information reduces the efficiency of visual foragers [20–23], and can have larger-scale ecological impacts, for example on species abundances and biodiversity [24–26]. At relatively high levels of turbidity (>50 Nephelometric Turbidity units, NTU), group cohesion has also been shown to be disrupted in fish shoals [27–29], although at lower levels, nearest neighbour distances can be maintained in turbid water [30,31], and in some cases, shoals can even be more cohesive at moderate turbidity compared to clear and highly turbid water [32]. These effects on group cohesion are consistent with the importance of vision in the shoaling behaviour of many species of fish [33–35], but also suggest some plasticity with robustness to moderate levels of turbidity.

Given the evidence that larger groups can show a greater capability in detecting and responding to stimuli, and that turbidity is an ecologically relevant environmental stressor that can be detrimental to both foraging and group cohesion in animals that rely on vision, we tested whether the effect of turbidity on foraging performance is dependent on group size.

Fish shoals of three different sizes were used (2, 6 and 10 individuals), each tested in clear and turbid water in a fully-factorial design. A recently-developed experimental protocol was adopted [12] which consists of an open arena (as often used in laboratory studies to quantify collective behaviour [33,36]) modified to allow a food stimulus to be presented at an unpredictable location to the test subjects with minimal disturbance (Fig 1). This allows for quantifying in high resolution the positions of the fish relative to the stimulus and to one another immediately before the stimulus appears, as well as their response to the stimulus.

**Fig 1.**
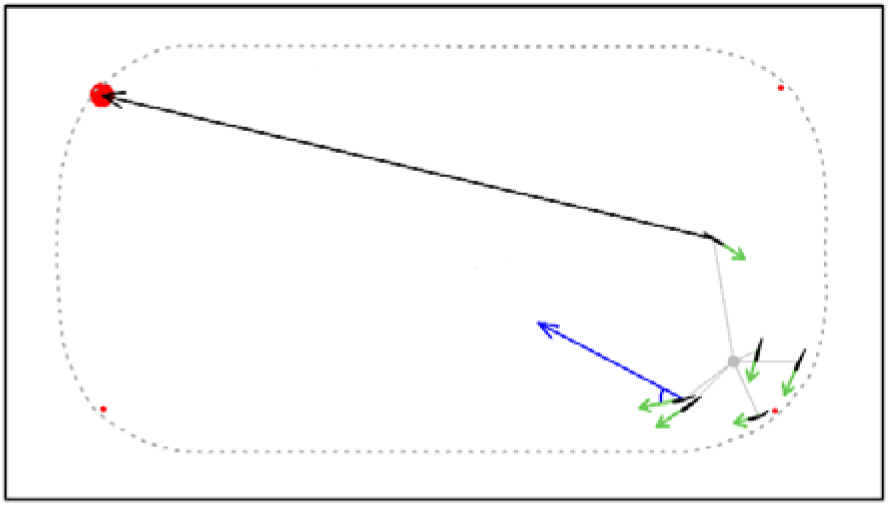
The test arena and the spatial and social variables used in the analysis. The figure is based on a thresholded still image from one of the experimental videos, thus the position of where the stimulus could appear (red circles) and the positions and size of the fish are to scale. The black arrow shows the distance from the stimulus (large red circle) to the nearest fish (i.e. shows the minimum distance to the stimulus). The minimum bearing to the stimulus is shown in blue, which is calculated as the angular difference between the orientation of the fish (small green arrow) and the direction from that fish toward the stimulus (blue arrow). The grey lines from each fish show the distance from each fish to the group centroid (grey circle), which is used to calculate the median distance to the group centroid, which is used as a measure of group cohesion. The orientation of each fish (small green arrows) is used to calculate the median group polarisation, which has a value of 1 when all fish face in the same direction, i.e. show high directional alignment, and 0 when the group is not polarised. The dotted grey line indicates the walls of the test arena.

## Results

We used shoals of three-spined sticklebacks (*Gasterosteus aculeatus*) as they are found in habitats ranging from clear, to periodically turbid, to chronically turbid [37]. We manipulated water clarity by conducting trials in either clear (<1 NTU) or moderately turbid water (∼10 NTU). A standardised stimulus was presented to the fish (a red-tipped dropping pipette) that elicits a foraging response without training, and also rewarded the fish with a single bloodworm when they came within two body lengths of the stimulus [12]. Either 682 or 697 presentations were included in the statistical analyses (see tables S1-S12 for details), with up to six presentations included from each of the 120 trials. Fish trajectories were automatically extracted using *idTracker* [38]. For many species, detecting and responding to important but unpredictable stimuli, such as food or a predator, is critical to survival, thus our main response variable as a measure of functional performance was the latency in time between the food stimulus appearing and the first attack on the stimulus or on the ejected bloodworm. The importance of explanatory variables affecting response variables was determined using a model comparison approach with the AICc (Akaike information criterion, corrected for small sample sizes) [39].

### The impact of water turbidity is independent of group size

As expected from extensive research in swarm (i.e. collective) intelligence in animal groups [7,8,40], larger shoals were faster in attacking the food stimulus, showing reduced latencies in both clear and turbid water (coefficient = -0.26, standard error (S.E.) = 0.027, 95% confidence intervals (C.I.) = -0.31 to -0.21; Fig 2A; table S1). Such a group size effect could be explained by larger groups being more likely to have some individuals closer to the stimulus and with the stimulus directly ahead of them, both which have been shown to improve responses when group size is held constant [12]. To account for these factors, we expanded the model to include as additional explanatory variables the distance of the nearest fish to the stimulus and the minimum angular bearing of the stimulus to a fish (Fig 1) just before the presentation. Both the minimum distance to and minimum bearing of the stimulus affected the latency to attack by making the model more likely when they were added (minimum distance: coefficient = 0.076, S.E. = 0.022, 95% C.I. = 0.034 to 0.12; minimum bearing: coefficient = 0.13, S.E. = 0.020, 95% C.I. = 0.092 to 0.17). The minimum bearing of the stimulus had a greater effect than the minimum distance to the stimulus, although group size also remained an important variable to include in the models (coefficient = -0.19, S.E. = 0.028, 95% C.I. = -0.25 to -0.14; table S2). Thus, geometric factors alone could not fully explain the effect of group size on the improved latency to attack.

**Fig 2.**
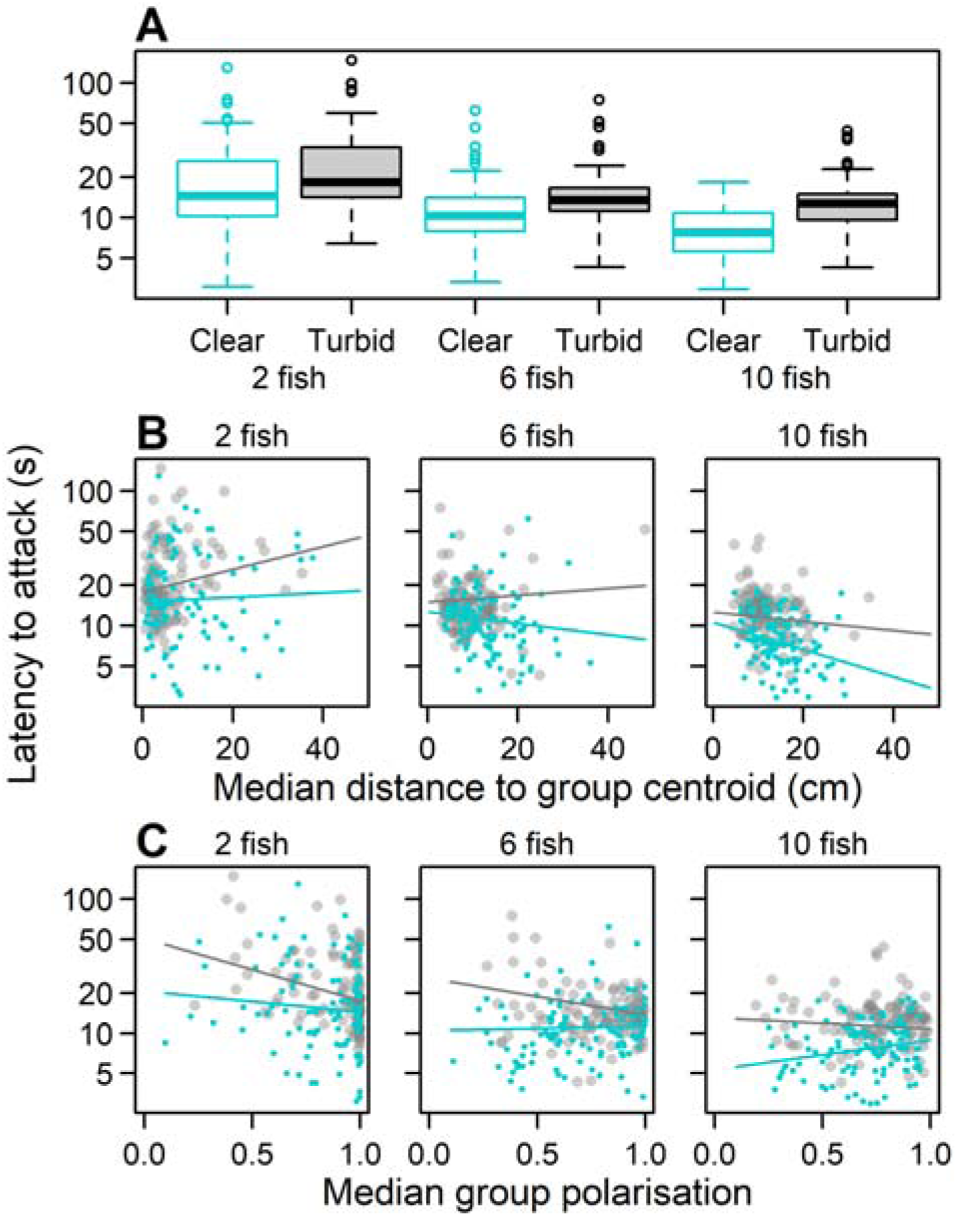
The effect on the latency to attack of water clarity and group size, cohesion and polarisation. Data from the clear water treatment is displayed in light blue, and from the turbid treatment in grey. In B, smaller values of distance to the group centroid are more cohesive groups. The boxplots in A show the medians as the thick horizontal lines, interquartile ranges are enclosed by the box, the most extreme data points within 1.5 × the interquartile range are shown by the whiskers, and empty circles show outliers beyond the whiskers. Lines of best fit in B and C are based on the LMM coefficients from the most likely models (tables S3 and S4, respectively).

Also consistent with predictions and previous work, the latency to attack was slower in turbid water (coefficient = 0.34, S.E. = 0.054, 95% C.I. = 0.23 to 0.45; table S1). However, there was no evidence for an interaction between water clarity and group size; the effect of group size was similar in both clear and turbid water, and latencies were consistently slower in turbid water regardless of group size (Fig 2A). Thus, the net effect of increased group size was neither to improve, nor diminish, resilience to reduced visual information from turbidity.

### Collective behaviour mediates the effects of turbidity and group size

Although there is a long history of exploring the effects of group size in studying animal sociality [40], more recently there is growing evidence that other properties of groups are also important for group functioning [8,41–43]. Based on the spatial positions and orientations of fish in the one second prior to the food stimulus appearing, we analysed whether the effect of turbidity and group size were mediated by the collective behaviour of the group. There was strong evidence from the model comparisons that including an interaction term between group cohesion (measured as the median distance to the group centroid; Fig 1) and water clarity, and between group cohesion and group size, improved the likelihood of the model (group cohesion × water clarity: coefficient = 0.11, S.E. = 0.043, 95% C.I. = 0.023 to 0.19; group cohesion × group size: coefficient = -0.076, S.E. = 0.022, 95% C.I. = -0.12 to -0.034; table S3). In turbid water, latencies in small groups were faster when cohesion was greater, with this effect being lost in larger groups in turbid water (Fig 2B). In contrast, cohesion did not affect the latencies of small groups in clear water, but being more cohesive was associated with slower latencies in larger groups in clear water (Fig 2B). Thus, the effect of cohesion on the latency to attack was dependent on water clarity, and was evident at different group sizes.

Group polarisation, a measure of coordination in orientation within animal groups (Fig 1), is another common metric used to quantify collective behaviour, and is known to vary between groups, populations and species [35,41,44] and to be functionally important [45,46]. As with group cohesion, polarisation during the one second prior to the food stimulus appearing interacted with both water clarity and group size (polarisation × water clarity: coefficient = -0.15, S.E. = 0.040, 95% C.I. = -0.23 to -0.067; polarisation × group size: coefficient = 0.073, S.E. = 0.021, 95% C.I. = 0.032 to 0.11; table S4). Consistent with a previous study using shoals of eight three-spined sticklebacks [12], in clear water the latency to attack was faster in more disordered, larger groups, i.e. those with lower polarisation (Fig 2C). However, in smaller groups in clear water, this effect weakened and was lost in pairs of fish. In contrast, groups of two in turbid water showed a negative association between polarisation and the latency to attack, i.e. more polarised groups were faster to attack, and this effect became weaker as group size increased. Together with the effect of group cohesion, these results reveal that collective behaviour is important in small groups in turbid water, but in larger groups in clear water.

Although there are often associations in fish shoals between polarisation and swimming speed, and cohesion and swimming speed [47,48], we found no evidence for an interaction term between water clarity and the speed of the fish in the one second prior to the food stimulus appearing on the latency to attack (table S5), suggesting that the effects of polarisation and cohesion in clear versus turbid water could not be explained by differences in speed. We also found no evidence that the median bearing of the nearest neighbour before the stimulus appeared, as a measure of how individuals are positioned relative to one another within a shoal, affected the latency to attack either as a main effect or interaction with other variables (table S6).

### Adaptive collective behaviour can improve responses to stimuli

If group cohesion and polarisation have effects on being able to respond to stimuli that differ between clear and turbid water, groups may adapt their cohesion and polarisation depending on environmental conditions before stimuli are present to maximise their ability to respond. Consistent with the effects on responses in clear and turbid water, in the one second immediately before the food stimulus appeared, the fish in turbid water were more cohesive (coefficient = -0.33, S.E. = 0.060, 95% C.I. = -0.45 to -0.21; table S7) and more polarised (coefficient = 0.32, S.E. = 0.086, 95% C.I. = 0.16 to 0.49; table S8) than fish in clear water (Fig 3; this was controlling for group size as a main effect, although there was some support for an interaction between group size and water clarity in predicting group cohesion (ΔAICc of the main-effects only model from the most likely model with the interaction = 1.8, table S7)). Together, these differences depending on water clarity suggest that conformity in spatial position and orientation are favoured in turbid water, while differentiation is favoured in clear water, and demonstrates that animal groups can adjust the balance between conformity and differentiation depending on environmental conditions [49].

**Fig 3.**
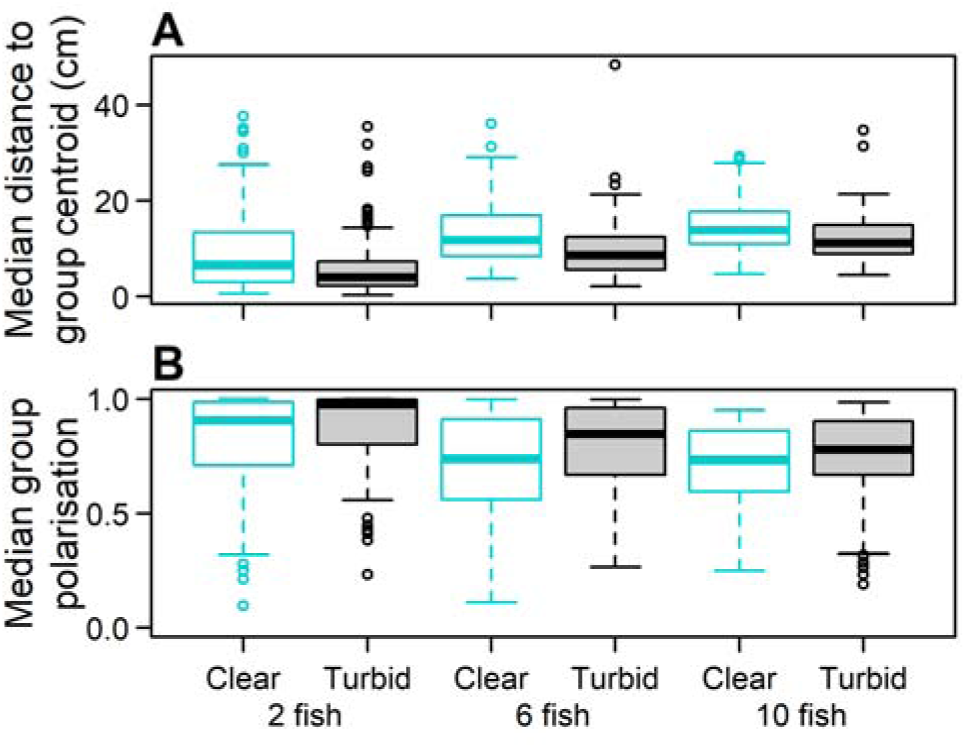
The effect of water clarity and group size on group cohesion and group polarisation. Plotting as in fig 2A.

From the most likely models of how water clarity and group size affect group cohesion and group polarisation (table S7, S8), and how group cohesion and group polarisation affect the latency of attack (table S3, S4), we can estimate the impact of the fish becoming more cohesive and polarised in turbid water on the latency to attack the stimulus. The magnitude of the effect of the observed increase in group cohesion was similar to that of group polarisation, where the change in collective behaviour in turbid water led to an estimated improvement in the latency to attack in groups of two fish of 1.0 second for group cohesion and, separately, a 0.8 second improvement due to group polarisation, representing a reduction in latencies of 4.6 and 4.0%, respectively (table 1). Corresponding to the reduced effect of cohesion and polarisation on the latency to attack in turbid water as group size increased (Fig 2B,C), this effect diminished as group size increased (table 1).

**Table 1.** The estimated change in the latency to attack the stimulus due to the fish increasing group cohesion and group polarisation in turbid water. The estimates are based on the most likely models in predicting group cohesion (measured as the median distance to the group centroid), group polarisation, and the latency to attack with group cohesion and group polarisation as predictors (tables S3, S4, S7, S8). The latency to attack if the fish did not change their behaviour in turbid water is based on their collective behaviour in clear water, with the latency then estimated for these values of group cohesion and polarisation if the fish were tested in turbid water. The distance to the group centroid is given in cm. The estimated improvement in the latency as a % is based on the change in latency compared to the latency in clear water.

| Collective behaviour (CB) | Group size | CB in clear water | CB in turbid water | Change in latency (s) | Change in latency (%) |
| --- | --- | --- | --- | --- | --- |
| Distance to group centroid | 2 | 8.7 | 6.3 | 1.0 | 4.6 |
| Distance to group centroid | 6 | 11.8 | 8.5 | 0.3 | 1.9 |
| Distance to group centroid | 10 | 15.8 | 11.4 | -0.4 | -3.5 |
| Group polarisation | 2 | 0.84 | 0.88 | 0.8 | 4.0 |
| Group polarisation | 6 | 0.76 | 0.82 | 0.5 | 3.3 |
| Group polarisation | 10 | 0.65 | 0.72 | 0.2 | 1.3 |

### Water turbidity hinders rapid collective detection of stimuli

While the distance of the nearest fish to the stimulus and the minimum bearing of the stimulus to a fish did not fully explain the effect of group size on the latency to attack, they both interacted with water clarity (minimum distance × water clarity: coefficient = -0.14, S.E. = 0.042, 95% C.I. = -0.23 to -0.063; minimum bearing × water clarity: coefficient = -0.098, S.E. = 0.039, 95% C.I. = -0.17 to -0.021; table S9, 10). In clear water, the latency to attack the stimulus was faster when the nearest fish was closer to the stimulus (Fig 4A). In turbid water, the distance to the nearest fish had no effect. Similarly, the minimum bearing of the stimulus to a fish was positively associated with the latency to attack in clear water, and less positively associated in turbid water (Fig 4B). These results are consistent with the fish often detecting the stimulus soon after it first appeared in clear water, so that the distance and bearing of the stimulus directly affected the latency to attack. In contrast, in turbid water the distance and bearing of the stimulus poorly predicting the latency to attack suggests that the fish were instead more likely to detect the stimulus after a longer delay, when the fish had moved and the distance and bearing of the stimulus had changed.

**Fig 4.**
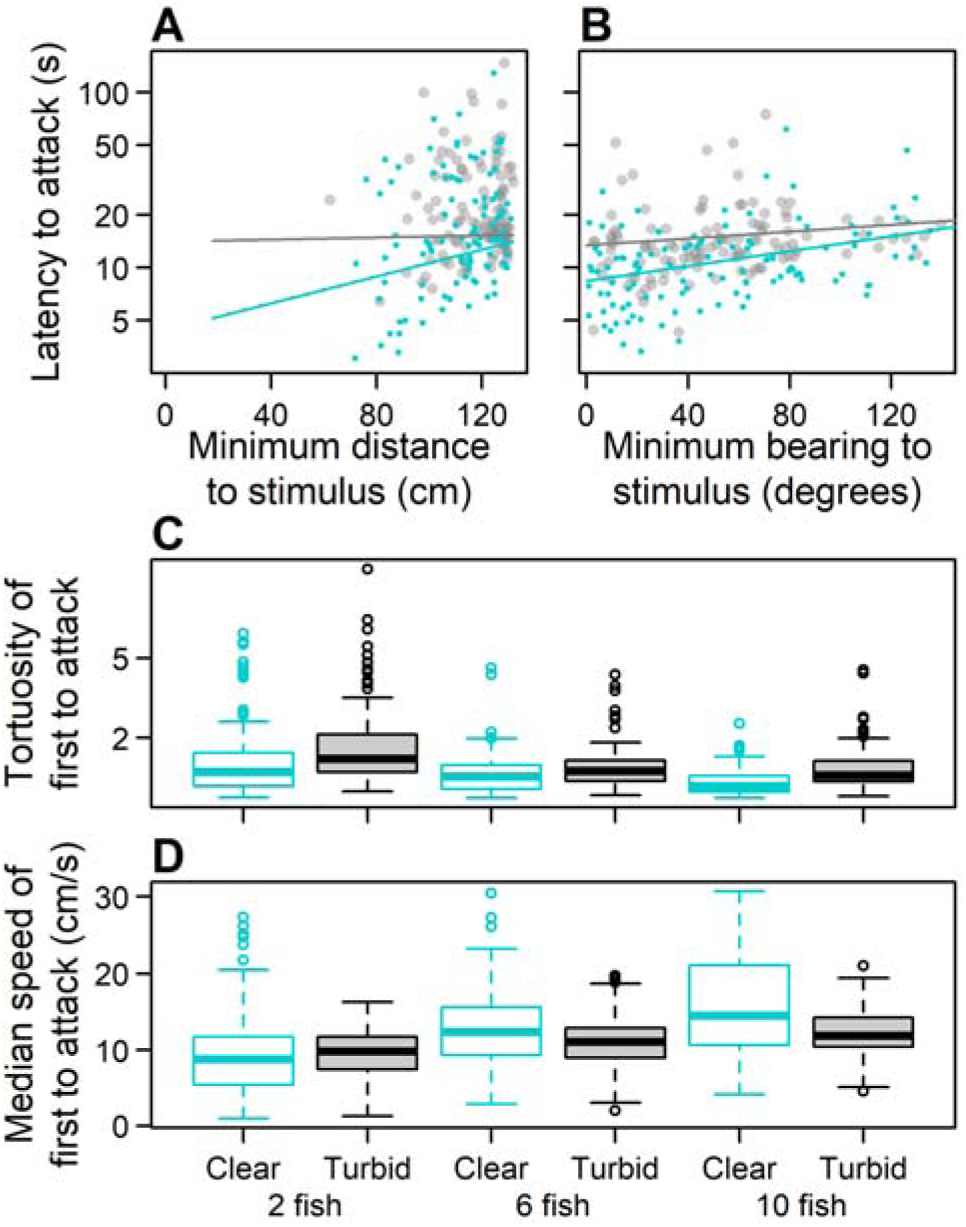
The effect of the minimum distance and the minimum bearing to the stimulus on the latency to attack, and the effect of water clarity and group size on the trajectory toward the stimulus of the fish to first attack. Data from the clear water treatment is displayed in light blue, and from the turbid treatment in grey. Lines of best fit in A and B are based on the LMM coefficients from models with the interaction between water clarity and the minimum distance to the stimulus (panel A, table S9) and between water clarity and the minimum bearing of the stimulus (panel B, table S10). The boxplots in C and D follow the same plotting as in fig 2.

To test this further, for the fish that made the first attack, we analysed its movement trajectory between the stimulus appearing and when it attacked the stimulus; sticklebacks respond after detecting food with direct, straight accelerated trajectories [12]. The fish trajectories were straighter (less tortuous) in clear water compared to turbid water (coefficient = 0.41, S.E. = 0.070, 95% C.I. = 0.27 to 0.55; Fig 4C, table S11), consistent with detection of the stimulus soon after it appeared in clear water and later in turbid water, and in agreement with previous analyses of the trajectories of foragers in turbid water [20]. In the larger groups tested, but not the smaller groups, the median swimming speed of these fish was also faster in clear water than in turbid water (group size × water clarity: coefficient = -17.78, S.E. = 6.053, 95% C.I. = -29.75 to -5.83; Fig 4D, table S12), also indicating that detection of the stimulus occurred earlier in clear water.

## Discussion

Studies of collective behaviour have focused on the mechanisms underlying group formation and maintenance, information transfer, and collective decision making [41,42,50]. Far fewer studies have linked collective behaviour directly to functionally important outcomes [12,46,51,52], or considered the impact on collective behaviour of altered environmental conditions from anthropogenic change [10,11]. Our results demonstrate the importance of collective behaviour in how environmental conditions affect the response of groups to ecologically-relevant stimuli. Moreover, we provide evidence that fish are phenotypically plastic in their collective behaviour by adjusting it over the short term, which improves the responses of small groups to stimuli when environmental conditions, such as turbidity, reduce visibility [53]. Such a short-term behavioural flexibility may mitigate some of the negative impacts of turbidity before longer-term sensory adaptation to turbid water during development and adulthood can take place [54,55]. While group size has repeatedly been shown to be important in improved foraging and reducing predation risk which can be driven by swarm (i.e. collective) intelligence [7,40], our results suggest that other group-level properties (cohesion and polarisation) are those that directly interact with turbidity in determining the response. Our results suggest that quantifying collective behaviour, as well as manipulating or measuring group size, would be a fruitful avenue for future research in collective intelligence and how social animals are adapting to, or susceptible to, environmental change [56,57].

A typical assumption in studies of collective behaviour is that the behaviours individuals use to interact with one another are ‘rules of interaction’ that do not vary over time or context, including environmental conditions [50]. Changes in collective behaviour due to altered environmental conditions are often considered as arising from constraints, for example reduced visibility in turbid water, or as by-products of physiological changes, for example increased swimming speed in warmer water [10]. Instead, animals may adjust how they react to their neighbours to maximise food intake or minimise predation risk in a manner that is adaptive depending on the context [58], and this is consistent with empirical work demonstrating that collective movement changes over different time scales, from short-term plastic behavioural responses to evolutionary change over generations [10,41]. Our findings that in moderately turbid water fish shoals are able to increase cohesion and polarisation, which has a functionally-important improvement in the speed at which they find unpredictable food in their environment (at least in small groups), is a clear demonstration of this phenomenon. This calls for simulation modelling of groups that can adaptively alter their interaction behaviours, with likely application in swarm robotics [59,60].

## Materials and Methods

### Experimental design

Our objective was to test whether larger groups were more or less susceptible to the negative impacts of water turbidity on the detection of unpredictable food resources [12], and whether this was affected by the collective behaviour of the group. We thus used a fully factorial design of testing shoals at three group sizes (2, 6 and 10 individuals), with each group size tested in clear and turbid water. Shoals of fish were filmed from above and their motion trajectories were tracked in two-dimensions using high resolution tracking software [38].

### Experimental subjects and housing

Three-spined sticklebacks were purchased from Carp Co. (www.carpco.co.uk) and transported by van to the University of Bristol in August 2021. Fish were housed in three glass tanks (70cm (L) × 40cm (W) × 36cm (H)), each containing approximately 70 individuals. Fish were fed daily with defrosted bloodworms and pellet feed. The laboratory was maintained at an air temperature of 16°C with a photoperiod of 11 : 13 hours (light : dark cycle). Fish were 3.9 ± 0.37 cm standard length (mean ± SD) at the time of testing. The temperature and light conditions stopped the fish from entering breeding condition, and females and males are not morphologically distinct when not breeding; fish were thus not sexed. After the study, fish were retained in the laboratory for use in other experiments.

### Experimental Setup

The test arena (136cm (L) x 73cm (W) x 50cm (H)) was constructed using 3mm white foamed polyvinyl chloride (PVC) sheets (Fig 1), following the design in [12]. The arena was filled with aged tap water to a depth of 8 cm. A hole (8 mm in diameter) was made in each of the four corners of the arena (Fig 1) at 6 cm above the arena floor. A Sony AX53 video camera was placed centrally above the arena at a height of 124 cm above the water surface, and set to film in 4K resolution at 25 frames per second. A black curtain was draped over and around the arena to avoid disturbance. To allow the experimenter to view the arena during the trials, a second video camera (Panasonic HC-X920) was also positioned above the arena and linked to a video monitor located on the other side of the curtain. Six pipettes were prepared by wrapping 2cm of red electrical PVC tape around the pipette tip.

### Experimental Protocol

Prior to the trials, 72 fish from the holding tanks were haphazardly selected as test fish and separated into two groups, with each group held in different holding tanks (70cm (L) × 40cm (W) × 36cm (H)). Trials were conducted between 10:00 and 16:00 for four days a week from Monday to Thursday for five weeks from 14th February to 17th March 2022. For each day of trials, we tested fish from one of the two holding tanks, alternating between tanks each day so that no fish was tested on any two consecutive days. Each individual fish was only tested once each day; after testing, they were transferred to a separate holding tank for the remainder of that day. Fish tested on a test day were fed only during the trials and at the end of the day after testing had been completed to standardise hunger at the start of each trial.

Each group size was tested once in the morning and once in the afternoon, totalling six trials per day, with the order of testing the different group sizes randomised within each morning and each afternoon (i.e. a complete random block design). For the water clarity treatment (turbid or clear), we conducted two consecutive days of turbid trials and two consecutive days of clear trials per week. Which treatment occurred first in each week was also randomised. Turbidity was increased by adding 1.2 grams of kaolin clay powder, which was suspended in arena water within a small container before being poured into the arena and stirred. For the trials with turbid water, prior to adding the fish, the arena water was stirred for 30 seconds to ensure the suspension of the clay. Immediately after stirring, two turbidity readings were taken from the centre of each half of the arena (11.7 ± 1.4, mean Nephelometric Turbidity Unit (NTU) ± SD) using a Thermo Scientific Orion AQUAfast AQ3010 turbidity meter. These turbidity measurements were repeated immediately after each turbid-water trial (9.26 ± 1.38, mean NTU ± SD). For days of testing the clear treatment, the same procedure was used to measure turbidity but only before each trial (0.73 ± 0.12, mean NTU ± SD) and after all trials on each test day (0.77 ± 0.09, mean NTU ± SD).

Before each trial, each of the six pipettes were loaded with a single defrosted bloodworm (chironomid larvae). At the start of each trial, the fish were placed into the centre of the arena and the video recording began. The fish were given two minutes to acclimatise to the arena before the first presentation was made. This involved gently inserting the tip of one of the pipettes into the hole in the corner of the arena that was furthest from the fish. If the fish were distributed across both halves of the arena, the presentation was delayed until all the fish had moved into one half of the arena. Once a fish had swam within two body lengths of the pipette tip, the bloodworm was ejected. Once the bloodworm had been eaten, 2 minutes were given before the next presentation, thus allowing time for the fish to resume their swimming behaviour; the mean time elapsed between the food item being consumed and the next presentation being made within a trial (± SD) was 2.23 ± 0.34 minutes. A total of six presentations were made per trial. In 21 presentations, the fish did not consume the ejected bloodworm; as this suggested that the fish were not motivated to feed, these presentations were removed from our analyses, with 699 presentations remaining.

### Ethics statement

The procedures using animals were approved by the University of Bristol Animal Welfare and Ethical Review Body (UIN UB/17/060). The test subjects were retained in the laboratory for use in future experiments.

### Video Processing and Data Extraction

The video of each trial was converted from 4K (.mp4) to 1920 × 1080 pixels (.m4v) using *Handbrake* 1.5.1. We used the automated two-dimensional animal tracking software *idTracker* 2.1 [38] to obtain the *x*, *y* coordinate positions for the centre-of-mass of each individual fish for each frame of the video. The frame at which each presentation was made (i.e. when the red pipette tip appeared) and when the food item or the red pipette was attacked (whichever occurred first) were manually recorded from the video; the number of frames that elapsed was converted into seconds and used as the latency to attack. The *x*, *y* coordinates of the red stimulus was also determined for each presentation. Due to errors in the automated tracking, 17 presentations were removed from the analysis involving variables calculated before the stimulus appeared, and 2 were removed from the analysis of variables after the stimulus appeared.

### Statistical Analysis

*R* 4.1.2 was used for all data processing and analysis. Based on the coordinates of the stimulus and of the fish in the frame the stimulus appeared, we calculated the Euclidean distance between each fish and the stimulus; the smallest value was used to calculate the minimum distance to the stimulus (Fig 1). With the additional coordinates on where each fish was in the preceding frame, the bearing of the stimulus from each fish was calculated using the cosine rule. The smallest value was used to calculate the minimum bearing to the stimulus. As a measure of group cohesion, we calculated the group centroid as the mean value of the *x* coordinates and mean value of the *y* coordinates for all the fish in the group.

The Euclidean distance from each fish to this centroid was calculated. The median value of all distances to the group centroid in the frames during the one second before the stimulus appeared was used to calculate the median distance to the group centroid. Group polarisation was calculated as detailed in [12] for each frame in the one second before the stimulus appeared; the median value was taken over these frames to give the median group polarisation. The Euclidean distance each fish moved per frame in the frames one second before the stimulus appeared were averaged using the median to give the median speed. For the fish to first attack the stimulus, the tortuosity of its trajectory between when the stimulus first appeared and when the attack was made was calculated as the distance travelled between these two points divided by the straight line distance between these two points. The fish’s median speed over this time was also calculated.

(Generalised) Linear Mixed Models, (G)LMMs, were used to analyse response variables of interest. Models differing in their fixed effects were constructed and compared using the AICc (Akaike information criterion corrected for small sample sizes). The *DHARMa* package 0.4.6 [61] was used to test whether the model assumptions were met in each set of model comparisons. Specifically, QQ plots were used to test whether the residuals were distributed as assumed by the error distribution specified in the model, residual versus fitted value plots were used to test whether there was any association between the residuals and the fitted values, and for the negative binomial model, the *DHARMa* package was used to test whether the dispersion parameter was approximately equal to one.

The time delay between the stimulus appearing and when either the stimulus or the bloodworm were first attacked by a fish was used as the latency to attack, and was the response variable in most of the models. This was natural log transformed to meet the assumptions of LMMs. The median distance to the group centroid was analysed using a negative binomial GLMM, the median group polarisation was analysed using a beta distributed GLMM, and the (natural log transformed) tortuosity of the first fish to attack was analysed using a Gamma distributed GLMM. The median speed of the fish after the stimulus appeared was analysed using an LMM; transforming the response variable was not required in this case.

The *lme4* 1.1-35.2 [62] package was used for all models except the beta and Gamma GLMMs where the *glmmTMB* 1.1.9 package [63] was used. Details of the fixed effects that varied between models in the model comparisons are given in the tables S1-S12. In all models we also included as fixed effects the presentation number within a trial and the testing day, as shoaling in three-spined sticklebacks is known to lessen over repeated testing [44]; we also factored in the trial number within a day to account for any diurnal effects. All continuous explanatory variables were scaled in the models (i.e. values had the mean value subtracted and were divided by the standard deviation). As there were multiple presentations per trial, trial number was included as a random effect, nested in the housing tank of the fish tested in that trial.

## Supporting information

Tables S1 to S12

## Author Contributions

Conceptualization: CCI, SAR, JC Funding acquisition: CCI Methodology: JC, CCI Resources: CCI

Investigation: JC, CCI Data curation: CCI, JC Formal analysis: CCI, SAR Visualization: CCI Supervision: CCI

Writing—original draft: CCI, SAR Writing—review & editing: CCI, SAR, JC

## Supporting Information

**Table S1.**
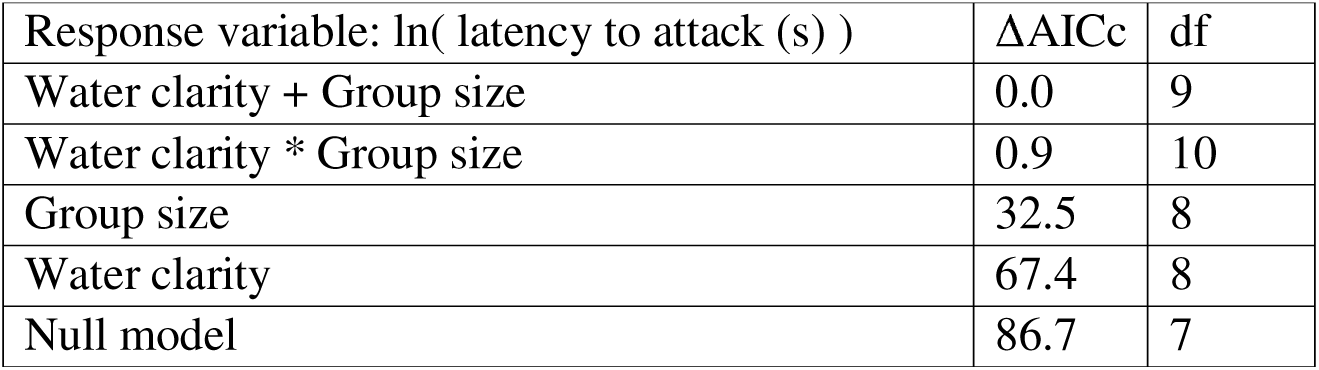
Effects of turbidity and group size on the latency to attack. Model comparison using the ΔAICc (difference in the Akaike information criterion, corrected for small sample sizes, between the model and the most likely model) to determine whether group size, water clarity (clear or turbid water) or the interaction between these variables had an effect on the latency to attack (natural logarithm transformed). The df is the number of parameters estimated in each model. N = 682 presentations.

**Table S2.** Effects of stimulus distance and stimulus bearing on the latency to attack. Model comparison using the ΔAICc to determine whether the effect of group size on the latency to attack could be explained by the minimum distance to and minimum bearing of the stimulus. N = 682 presentations.

| Response variable: ln( latency to attack (s) ) | $\Delta AIC_c$ | df |
| --- | --- | --- |
| Water clarity + Group size + Min. distance to stimulus + Min. bearing to stimulus | 0.0 | 11 |
| Water clarity + Group size + Min. bearing to stimulus | 10.1 | 10 |
| Water clarity + Group size + Min. distance to stimulus | 39.7 | 10 |
| Water clarity + Min. distance to stimulus + Min. bearing to stimulus | 41.3 | 10 |
| Water clarity + Group size | 53.3 | 9 |
| Water clarity + Min. bearing to stimulus | 63.6 | 9 |
| Water clarity + Min. distance to stimulus | 92.5 | 9 |

**Table S3.** Effects of group cohesion on the latency to attack. Model comparison using the ΔAICc to test for interactions on the latency to attack between group cohesion (median distance to the group centroid) before the stimulus appeared, water clarity and group size. N = 682 presentations.

| Response variable: ln( latency to attack (s) ) | $\Delta AICc$ | df |
| --- | --- | --- |
| Water clarity * Median distance to group centroid +<br>Group size * Median distance to group centroid | 0.0 | 12 |
| Water clarity * Group size * Median distance to group<br>centroid | 4.1 | 14 |
| Group size * Median distance to group centroid + Water<br>clarity | 4.1 | 11 |
| Water clarity * Median distance to group centroid + Group<br>size | 10.2 | 11 |
| Water clarity + Group size | 11.6 | 9 |
| Water clarity + Group size + Median distance to group<br>centroid | 13.7 | 10 |

**Table S4.** Effects of group polarization on the latency to attack. Model comparison using the ΔAICc to test for interactions on the latency to attack between group polarization before the stimulus appeared, water clarity and group size. N = 682 presentations.

| Response variable: ln( latency to attack (s) ) | $\Delta AICc$ | df |
| --- | --- | --- |
| Water clarity * Median group polarization +<br>Group size * Median group polarization | 0.0 | 12 |
| Water clarity * Group size * Median group polarization | 3.8 | 14 |
| Water clarity * Median group polarization + Group size | 9.9 | 11 |
| Group size * Median group polarization + Water clarity | 10.9 | 11 |
| Water clarity + Group size + Median group polarization | 20.5 | 10 |
| Water clarity + Group size | 23.3 | 9 |

**Table S5.** Effects of fish speed on the latency to attack. Model comparison using the ΔAICc to test for interactions on the latency to attack between the median speed of the fish before the stimulus appeared, water clarity and group size. N = 682 presentations.

| Response variable: ln( latency to attack (s) ) | $\Delta\text{AICc}$ | df |
| --- | --- | --- |
| Group size * Median speed + Water clarity | 0.0 | 11 |
| Water clarity * Median speed +<br>Group size * Median speed | 0.1 | 12 |
| Water clarity * Group size * Median speed | 2.4 | 14 |
| Water clarity + Group size + Median speed | 3.7 | 10 |
| Water clarity * Median speed + Group size | 3.9 | 11 |
| Water clarity + Group size | 53.2 | 9 |

**Table S6.** Effects of the bearing of the nearest neighbor on the latency to attack. Model comparison using the ΔAICc to test for interactions on the latency to attack between the median bearing of the nearest neighbor (NN) before the stimulus appeared, water clarity and group size. N = 682 presentations.

| Response variable: ln( latency to attack (s) ) | $\Delta\text{AICc}$ | df |
| --- | --- | --- |
| Water clarity * Median bearing of NN + Group size | 0.0 | 11 |
| Water clarity + Group size | 0.9 | 9 |
| Water clarity * Median bearing of NN +<br>Group size * Median bearing of NN | 1.5 | 12 |
| Water clarity + Group size + Median bearing of NN | 1.8 | 10 |
| Group size * Median bearing of NN + Water clarity | 3.4 | 11 |
| Water clarity * Group size * Median speed | 4.0 | 14 |

**Table S7.** Effects of turbidity and group size on group cohesion. Model comparison using the ΔAICc to test whether group cohesion (measured as the median distance to group centroid) was affected by water clarity, group size, or the interaction between these variables. N = 682 presentations.

| Response variable: median distance to group centroid | $\Delta\text{AICc}$ | df |
| --- | --- | --- |
| Water clarity * Group size | 0 | 10 |
| Water clarity + Group size | 1.8 | 9 |
| Group size | 26.8 | 8 |
| Water clarity | 52.3 | 8 |
| Null model | 69 | 7 |

**Table S8.** Effects of turbidity and group size on group polarization. Model comparison using the ΔAICc to test whether group polarization was affected by water clarity, group size, or the interaction between these variables. N = 682 presentations.

| Response variable: median group polarization | $\Delta\text{AICc}$ | df |
| --- | --- | --- |
| Water clarity + Group size | 0.0 | 9 |
| Water clarity * Group size | 1.5 | 10 |
| Group size | 11.1 | 8 |
| Water clarity | 67.3 | 8 |
| Null model | 72.4 | 7 |

**Table S9.** Effects of interactions with stimulus distance on the latency to attack. Model comparison using the ΔAICc to test for interactions between the minimum distance to the stimulus just before the stimulus appeared, water clarity and group size. N = 682 presentations.

| Response variable: $\ln(\text{ latency to attack (s) })$ | $\Delta\text{AICc}$ | df |
| --- | --- | --- |
| Water clarity * Minimum distance to the stimulus + Group size | 0.0 | 11 |
| Water clarity * Minimum distance to the stimulus +<br>Group size * Minimum distance to the stimulus | 1.0 | 12 |
| Water clarity * Minimum distance to the stimulus * Group size | 5.0 | 14 |
| Water clarity + Minimum distance to the stimulus + Group size | 9.8 | 10 |
| Group size * Minimum distance to the stimulus + Water clarity | 10.7 | 11 |
| Water clarity + Group size | 23.4 | 9 |

**Table S10.** Effects of interactions with stimulus bearing on the latency to attack. Model comparison using the ΔAICc to test for interactions between the minimum bearing of the stimulus to a fish just before the stimulus appeared, water clarity and group size. N = 682 presentations.

| Response variable: ln( latency to attack (s) ) | $\Delta AICc$ | df |
| --- | --- | --- |
| Water clarity * Minimum bearing of the stimulus + Group size | 0 | 11 |
| Water clarity * Minimum bearing of the stimulus +<br>Group size * Minimum bearing of the stimulus | 1.5 | 12 |
| Water clarity + Minimum bearing of the stimulus + Group size | 4.1 | 10 |
| Water clarity * Minimum bearing of the stimulus * Group size | 4.7 | 14 |
| Group size * Minimum bearing of the stimulus + Water clarity | 5.8 | 11 |
| Water clarity + Group size | 47.2 | 9 |

**Table S11.** Effects of turbidity and group size on swimming tortuosity. Model comparison using the ΔAICc to test whether the trajectory of the fish that first attacked the stimulus or bloodworm was more or less tortuous in clear versus turbid water. N = 697 presentations.

| Response variable: ln( tortuosity of first fish to attack ) | $\Delta AICc$ | df |
| --- | --- | --- |
| Water clarity + Group size | 0.0 | 9 |
| Water clarity * Group size | 0.1 | 10 |
| Group size | 29.1 | 8 |
| Water clarity | 52.4 | 8 |
| Null model | 71.7 | 7 |

**Table S12.** Effects of turbidity and group size on swimming speed. Model comparison using the ΔAICc to test whether the trajectory of the fish that first attacked the stimulus or bloodworm was faster or slower in clear versus turbid water. N = 697 presentations.

| Response variable: median speed of first fish to attack | $\Delta AICc$ | df |
| --- | --- | --- |
| Water clarity * Group size | 0.0 | 10 |
| Water clarity + Group size | 6.3 | 9 |
| Group size | 16.7 | 8 |
| Water clarity | 47.2 | 8 |
| Null model | 53.9 | 7 |

